# The Origin of Cognitive Modules for Face Processing: A Computational Evolutionary Perspective

**DOI:** 10.1101/2024.07.18.604211

**Authors:** Jirui Liu, Xuena Wang, Jia Liu

**Affiliations:** Department of Psychological and Cognitive Sciences & Tsinghua Laboratory of Brain and Intelligence, Tsinghua University

**Keywords:** Cognitive Modules, Meta-Learning, Evolutionary Algorithm, Connectivity, Fusiform Face Area, face recognition

## Abstract

Despite extensive research, understanding how cognitive modules emerge remains elusive due to the complex interplay of genetic, developmental, and environmental factors. Computational modeling, however, provides a means of exploring their origins by directly manipulating these factors. Here we aimed to investigate the emergence of cognitive modules by developing the Dual-Task Meta-Learning Partitioned (DAMP) model, whose plastic architecture facilitates automatic structure optimization through a genetic algorithm that emulates natural selection by iteratively selecting for efficient learning fitness. We found that a specialized module for face identification robustly emerged in the DAMP model. Critically, the emergence of the face module was not influenced by the demands of cognitive tasks (identification versus categorization) or the type of stimuli (faces versus non-face objects). Instead, it was determined by the structural constraint of sparse connectivity within the network, suggesting that the face module may arise as an adaptation strategy to challenges posed by sparse connections in neural networks, rather than being an information processor required by certain stimuli or tasks. These findings provide a new evolutionary perspective on the formation of cognitive modules in the human brain, highlighting the pivotal role of the structural properties of neural networks in shaping their cognitive functionality.

## Introduction

The human brain demonstrates a profound capacity for specialized processing, exemplified by cognitive modules where distinct neural substrates specialize in processing different types of information (Fodor, 1983). These cognitive modules are hypothesized to result from evolutionary pressures that favor neural efficiency in processing specific kinds of information. This functional specialization is particularly evident in the human ventral temporal cortex (VTC), which contains cortical regions dedicated to processing various categories of objects (Grill-Spector & Weiner, 2014). A prime example is the fusiform face area (FFA), extensively studied as a cognitive module specialized for face recognition (Kanwisher, 2000; Kanwisher et al., 1997; Kanwisher & Yovel, 2006). This specialization is thought to be driven by the unique demands of recognizing faces at the individual level, crucial for social interactions and communications. Moreover, its reliable detection via functional magnetic resonance imaging (fMRI) has facilitated numerous studies exploring its functions and characteristics (e.g., X. Chen et al., 2023; Grill-Spector et al., 2004; Rossion et al., 2012; Tarr & Gauthier, 2000; J. Zhang et al., 2021). However, neuroimaging and neurophysiological studies on the origin of the face module present significant challenges due to the complex interplay of innate predispositions (e.g., Wilkinson et al., 2014; Zhu et al., 2010), developmental plasticity (e.g., Patriquin et al., 2016; Peelen et al., 2009; Tian et al., 2021), social interactions (e.g., Jack & Schyns, 2015; Todorov et al., 2015; X. Wang et al., 2022), and visual expertise (e.g., Gauthier et al., 2000; Zhao et al., 2022).

Recently, computational modeling with artificial neural networks (ANNs) has emerged as a promising alternative, allowing researchers to precisely control and manipulate potential influencing factors (e.g., Kanwisher, Khosla, et al., 2023; M. Khosla & Wehbe, 2022; Lange et al., 2022; Richards et al., 2019; Sadeghnejad et al., 2024; Van Dyck & Gruber, 2023; Xue et al., 2024). This approach enables the detailed examination of how neural mechanisms might evolve and adapt under various conditions. One significant application of this approach is in the study of face recognition. Traditional ANNs designed for face recognition have demonstrated the emergence of face-selective units, validating their effectiveness (e.g., Blauch et al., 2022; Dobs et al., 2022). Beyond these specialized networks, randomly initialized ANNs have also shown the spontaneous development of face-selective units (Baek et al., 2021). Additionally, ANNs trained on tasks prioritizing spatial correlation costs, even without an explicit focus on face recognition, have exhibited face-selective responses (Lee et al., 2020). This phenomenon is also observed in domain-general ANNs without specific tuning for face recognition (Blauch et al., 2022) and even in unsupervised learning contexts (Keller et al., 2021). Remarkably, these face-selective units can develop in ANNs even without face experience (Xu et al., 2021). In addition to face recognition, ANNs tasked with fine-grained discrimination have shown functional specialization for non-face stimuli, such as cars and food (Dobs et al., 2022; Kanwisher, Gupta, et al., 2023). These findings from computational modeling suggest that neural mechanisms underlying functional specialization may be more adaptable than previously thought. However, a significant limitation of these studies is their reliance on pre-designed architectures, which do not accurately reflect the spontaneous emergence of specialized structures and functions seen in biological evolution. Accordingly, more advanced models are needed to better simulate the complex and dynamic nature of biological neural development.

Meta-learning, known as “learning to learn”, enables ANNs to adapt their learning processes and architectures based on new experiences or environments, rather than merely performing specific tasks (Hospedales et al., 2021). This capacity is crucial for developing more flexible and generalizable neural networks, as it allows them to modify their architectures in response to changing conditions. Here we employed a neural network that utilizes meta-learning with flexible neuron blocks (Fig. 1A), termed the Dual-Task Meta-Learning Partitioned (DAMP) model, and designed to evolve autonomously. The DAMP model features a genetic evolutionary meta-optimizer and three distinct types of neuron blocks: module 1 (M1), distributed (Dist), and module 2 (M2) blocks (Fig. 1A). Each block processes a specific type of information independently, thereby minimizing interference and enhancing processing efficiency.

**Figure 1:**
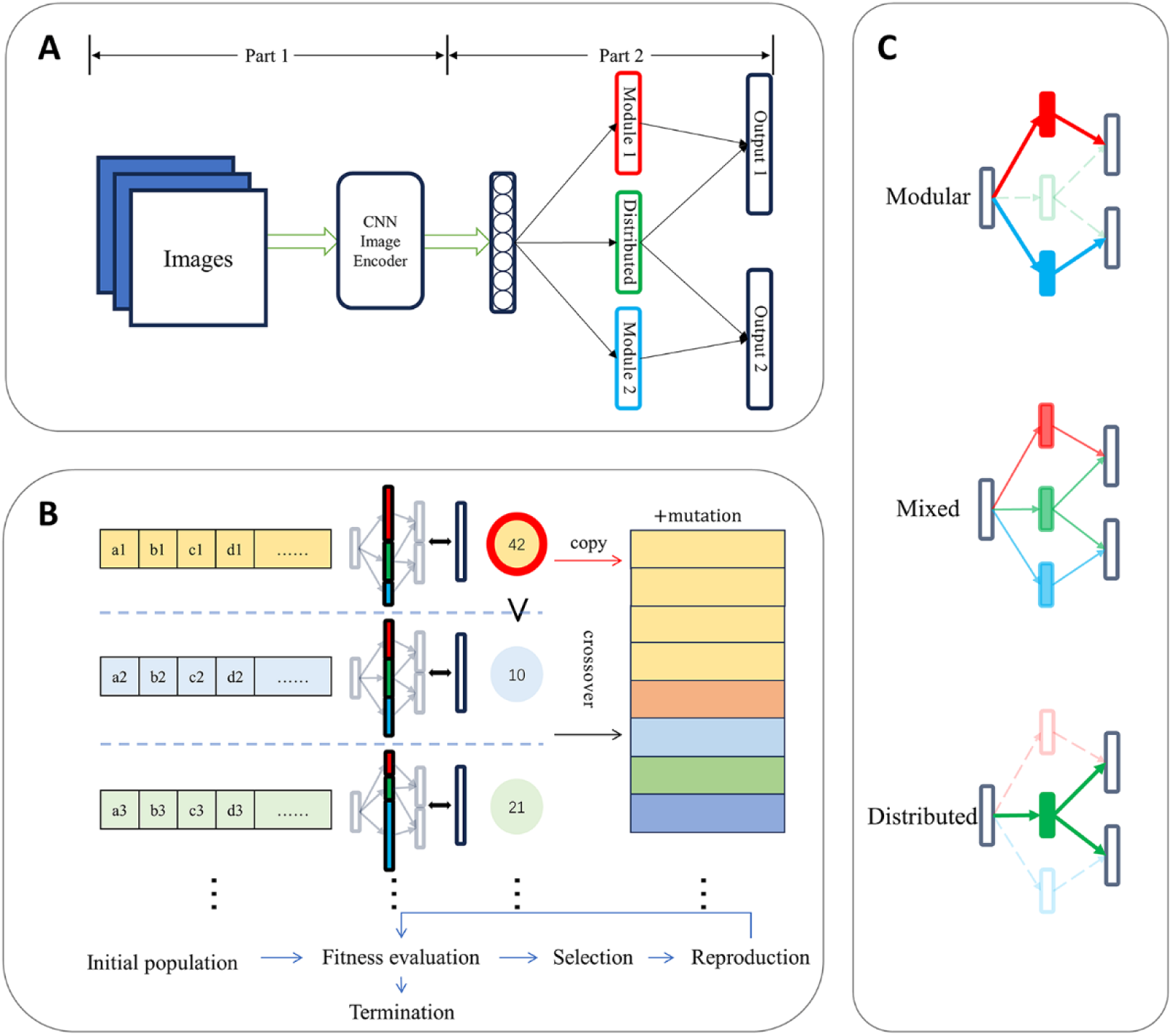
Meta-learning through evolutionary algorithms. (A) The DAMP model includes a pre- trained deep convolutional neural network (DCNN) encoder that translates input images into vector representations. These vectors are sent to a fully connected hidden layer, which then connects to three hidden neuron blocks: module 1 (M1), distributed (Dist), and module 2 (M2) blocks, each containing a variable number of neurons. These hidden neuron blocks are linked to two output neuron blocks: Output1 and Output2, each designated to fulfill a specific task (e.g., face identification and object categorization). (B) The evolutionary algorithm initiates by generating a gene population that determines the model’s hidden structure. This population is evaluated based on efficient learning fitness. The elite gene population with the highest fitness is selected, followed by recombination processes such as copying, crossover, and mutation. This selection process is iterated until the elite gene population stabilizes, indicating an optimal architecture has been achieved. (C) After the evolutionary process, the number of neurons in each hidden block defines three possible types of architectures: modular network (top), mixed network (middle), and distributed network (bottom). Solid rectangles indicate a neuron block with more than 0 neurons, while empty rectangles indicate a neuron block with 0 neurons.

The meta-optimizer utilizes an evolutionary algorithm to iteratively improve the model by simulating evolutionary processes, selecting for architectures that perform well across different tasks. Specifically, a gene population represents the network and learning parameters, which are evaluated in each generation to identify the highest fitness. New populations are generated from the selected high-fitness genes through processes of copying, crossover, and mutation, continuing until a steady state is achieved with a dominant elite gene (Fig. 1B). In this steady state, one gene consistently outperforms others across successive generations. The number of neurons in each block at this point indicates the degree of modularity, leading to three candidate architectures: modular, distributed, and mixed (Fig. 1C). Notably, this model does not require pre- defined architectures or manual hyperparameter tuning, allowing architectures to emerge naturally without any prior hypotheses. Similar models have been employed previously to investigate modularity in simplified what-where tasks (Bullinaria, 2007; Rueckl et al., 1989), demonstrating the potential of the DAMP model to provide insights into the spontaneous emergence of specialized structures and functions for face recognition, mimicking the processes of biological evolution and natural emergence of the face module.

In addition to developing the DAMP model, we tested two types of tasks: identification of exemplars at the individual level (i.e., recognizing specific instances within a category) and categorization at the basic level (i.e., classifying objects into general categories). These tasks are thought to be crucial in the formation of the face module, as they reflect different demands of recognizing faces and other objects (Gauthier et al., 2000; James & James, 2013; Rhodes et al., 2004). Non-face objects (e.g., cars, food, dogs, and daily objects) were also included to test the specificity of the DAMP model in forming specialized modules for faces versus other object types. This test allows us to assess whether the DAMP model selectively develops specialized neural substrates for face recognition or if similar processes occur for other objects, providing insight into the mechanisms of neural specialization. The aim of this study is to observe the natural emergence of specialized modules without predefined structures, providing a deeper understanding of how the face module and other specialized structures develop.

## Results

### The DAMP model

To more closely mimic evolutionary progress, we employed the DAMP model, which consists of two main components (Fig. 1A). The first component is a pre-trained DCNN encoder, based on a modified AlexNet (Krizhevsky et al., 2017) without the final fully connected layer. This modification allows for the extraction of comprehensive image features while reducing dimensionality to represent 80% of the explained variance, avoiding the constraints of the 1000-category representations typical in ImageNet. The second component comprises three hidden blocks: M1, Dist, and M2, which are fully connected to the output of the encoder. The total number of neurons (𝑁_𝑡𝑜𝑡𝑎𝑙_) across these blocks remains constant to control for the effect of neuron quantity, while the number of neurons in each block can vary. This adaptable architecture is particularly suitable for meta-learning, allowing the number of neurons in each block to adjust to different tasks under various evolving targets (i.e., meta-objectives). The model’s output is structured in two output layers (Output1 and Output2) for a dual-task setting, each receiving concatenated inputs from specific combinations of the blocks. For example, Output1 can be linked solely to M1 or receive input from both M1 and Dist. This design enables the model to encapsulate visual information relevant to each task in separate or shared blocks, thereby reflecting various degrees of modularity (Fig. 1C).

To quantify the degree of modularity, we introduced a modularity score ranging from 0 (fully distributed structure) to 1 (fully modular structure). The modularity score is calculated as 1 − (𝑁_𝐷𝐷𝐷𝑠𝑡_/𝑁_𝑡𝑜𝑡𝑎𝑙_), where 𝑁_𝐷𝐷𝐷𝑠𝑡_ is the number of neurons in Dist.

That is, a score of 1 indicates that no neurons are present in Dist after the evolutionary process, meaning Output1 and Output2 receive input exclusively from M1 and M2, respectively. This scenario suggests that the information from one block (e.g., M1) is sufficient for a specific task (e.g., face identification), thereby serving as a cognitive module (e.g., face module).

A genetic evolutionary algorithm is used as the meta-optimizer in meta-learning to optimize the architecture of the DAMP model, where the meta-objective is learning efficiency. This algorithm operates over multiple generations, with a set of 𝑁 genotypes serving as candidate solutions at generation 𝑡: 𝑃(𝑡) = {𝑥_1_, 𝑥_2_, …, 𝑥_𝑁_}. Each individual determining the network architecture is represented as a vector of genes:𝑥_𝐷𝐷_ = (𝑔_1_, 𝑔_2_, …, 𝑔_𝑘_). Each gene is a non-negative number representing a parameter of the network architecture (e.g., the number of hidden block neurons) or learning functions (e.g., learning rates and weight/bias initialization intervals). The evolutionary process includes two main steps in each generation (Fig. 1B): evaluating the fitness of candidate solutions and identifying the elite one: 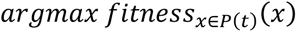. New populations 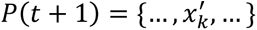 are reproduced through three mechanisms: copying (𝑥_𝑘_ = 𝑥_𝑒𝑙𝐷𝐷𝑡𝑒_(𝑡)), crossover (𝑥_𝑘_ =𝐶𝑟𝑜𝑠𝑠𝑜𝑣𝑒𝑟(𝑥_𝐷𝐷_, 𝑥_𝑗𝑗_)), and mutations 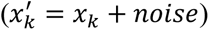 These mechanisms permute the architecture parameters to explore new potential solutions. Fitness serves as the direction of evolution to promote efficient learning, evaluated either by achieving a certain training accuracy (80%) faster in the early epoch or by attaining higher accuracy within a limited number of epochs (𝑒𝑝𝑜𝑐ℎ_𝑙𝐷𝐷𝑚_ = 50). The fitness function is defined as:

𝑓𝑓𝑓𝑓𝑡𝑛𝑒𝑠𝑠 = 𝑒𝑝𝑜𝑐ℎ_𝑙𝐷𝐷𝑚_ − 𝑒𝑝𝑜𝑐ℎ + 𝑎𝑐𝑐.

Since the total number of hidden neurons is constant (𝑁_𝑡𝑜𝑡𝑎𝑙_ = 𝑁_𝑀1_ + 𝑁_𝐷𝐷𝐷𝑠𝑡_ + 𝑁_𝑀2_), the three blocks compete for a larger number of neurons. Each generation yields one elite genotype (𝑥_𝑒𝑙𝐷𝐷𝑡𝑒_), and evolution stops when the elite network architecture remains consistent compared to the previous generation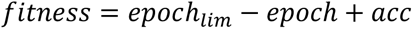 for network architecture gene This consistency indicates that a stabilized optimal solution has been reached: 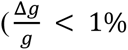), suggesting that its corresponding structure dominates others. Once evolution ends, we calculate the modularity score to quantify the degree of specialization within the network.

Neuron connections in brains are partially connected rather than fully connected to reduce volume and help maintain efficient neural processing while minimizing metabolic costs (Bullmore & Sporns, 2012). To decrease the connectivity level from the maximum value 1 (i.e., fully connected, completely dense coding), we can either reduce the connections between neurons in the blocks and those in the output layers or reduce the number of neurons in Dist (𝑁_𝐷𝐷𝐷𝑠𝑡_). That is, the DAMP model can achieve this by either randomly removing a portion of connections while treating all blocks equally (𝑓𝑓_𝐻𝐻𝐻_), or by specifically removing connections of the neurons in Dist that provide outputs to both output layers. Accordingly, the connectivity level (𝑓𝑓) is defined as:

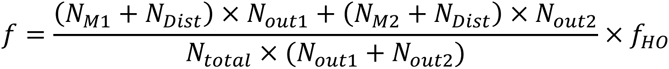

Where 𝑁_𝑀1_, 𝑁_𝑀2_, and 𝑁_𝐷𝐷𝐷𝑠𝑡_ represent the number of neurons in the respective blocks, and 𝑁_𝑜𝑢𝑡1_ and 𝑁_𝑜𝑢𝑡2_ represent the number of output neurons for the two tasks.

When the connectivity level is preset, the goal of the evolution is to adjust the pattern of connectivity between the blocks and the output layers to maximize learning efficiency. That is, if Dist benefits both tasks, its connections to the output layers should be maintained. However, if tasks are handled more efficiently by M1 and M2 alone, the number of neurons in Dist (𝑁_𝐷𝐷𝐷𝑠𝑡_) is reduced. In the first experiment, we set the connectivity level to 0.5, based on previous studies (Bullinaria, 2007), to explore how the DAMP model adapts its connectivity patterns to optimize learning efficiency while potentially forming the face module. By analyzing the resulting architectures, we can gain insights into the evolutionary mechanisms that drive the development of cognitive modules and their efficiency in processing specific tasks.

### Face module emerged in the DAMP model

We used a dual task of face identification and object categorization (i.e., the Face- Object dual task) to examine whether the face module emerges in the DAMP model. We selected face images from approximately 100 individuals (100 face exemplars per individual) and object images from approximately 100 object categories (e.g., airplanes, clocks; 100 exemplar images per category) from the VGGFace2 (Cao et al., 2018) and ILSVRC-2012 (Deng et al., 2009) databases (Fig. 2A). The total number of neurons (𝑁_𝑡𝑜𝑡𝑎𝑙_) was set to 900, which was equally assigned to three blocks (i.e., 300 neurons per block). Changes in the number of neurons in each block of 𝑥_𝑒𝑙𝐷𝐷𝑡𝑒_ at each generation, relative to 𝑁_𝑡𝑜𝑡𝑎𝑙_, are illustrated in Fig. 2B. These changes indicate how the evolutionary process adjusts the network’s architecture to enhance learning efficiency and potentially form specialized modules.

**Figure 2:**
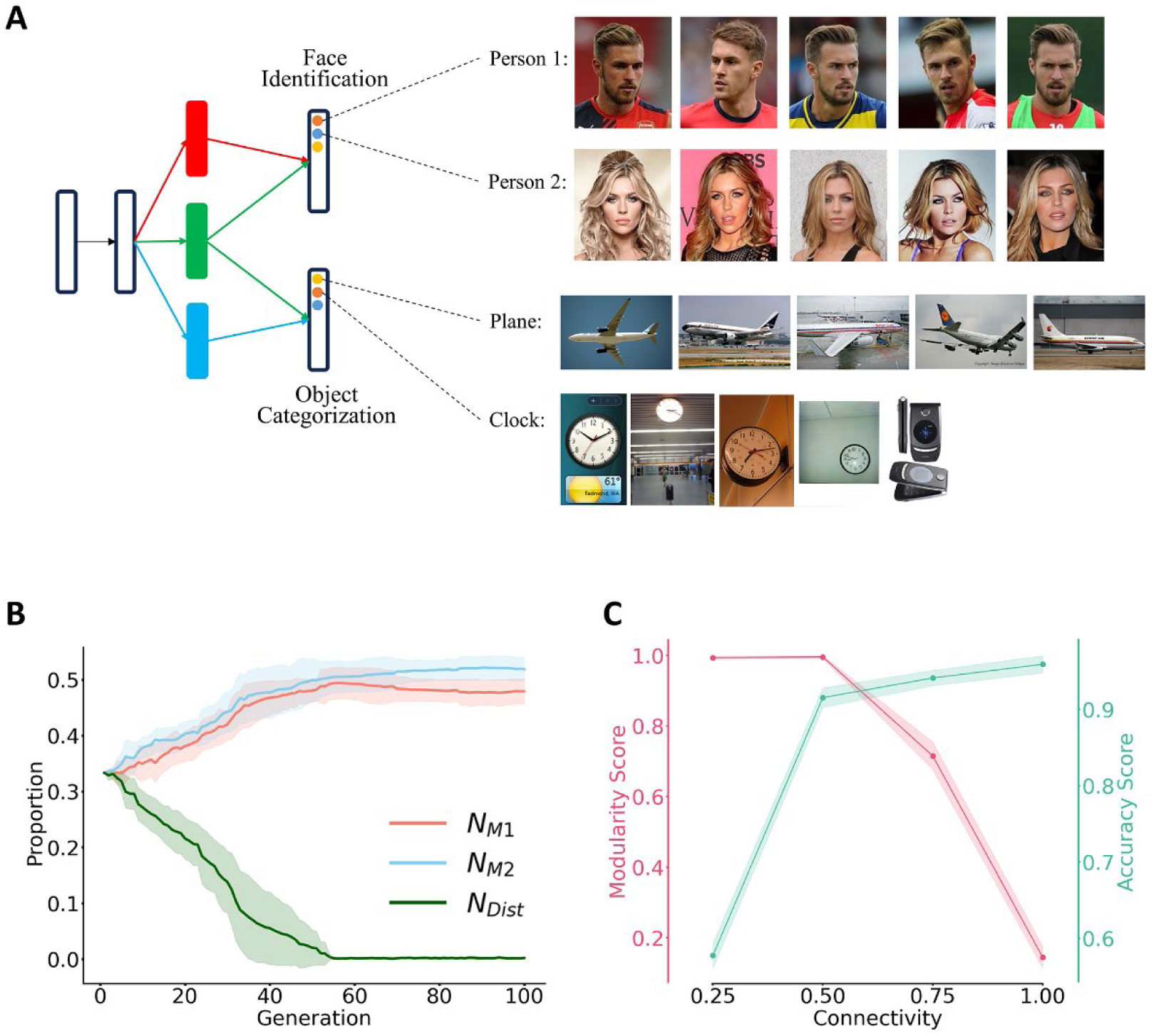
The emergence of the face module under the Face-Object dual task. (A) In the Face- Object dual task, each neuron of the output layer represents one individual (i.e., face identification, Output1) or one object category (i.e., object categorization, Output2). Examples of two individuals and two object categories (plane and clock) are shown. (B) Evolutionary dynamics of the neuron distribution among the blocks (M1: red, Dist: green, M2: blue). The x-axis denotes evolutionary generations, while the y-axis shows the number of neurons relative to 𝑁_𝑡𝑜𝑡𝑎𝑙_ . The solid line represents the mean changes, and the shaded area represents the standard deviation across 8 repeats.

Overall, the number of neurons in Dist (𝑁_𝐷𝐷𝐷𝑠𝑡_) decreased monotonically and approached zero after about 55 generations, indicating that the shared information carried by Dist became less important during evolution. In contrast, 𝑁_𝑀1_ and 𝑁_𝑀2_ increased concurrently, ultimately forming a modular structure. This shift reflects the model’s adaptation towards specialized processing units, enhancing learning efficiency for the Face-Object dual task. At the end of the evolution, the accuracy of the dual task was above 60% (chance level = 0.1%), and the modularity score was close to 1 (mean = 0.9992, SD = 0.0009), signifying a highly specialized and efficient structure. Specifically, M1 evolved into the face module, as Output1, designated for face identification, exclusively receives input from M1. Thus, a face module emerged in the DAMP model under the dual task of face identification and object categorization, illustrating the model’s ability to develop specialized modules for distinct tasks.

To examine the robustness of this finding, we further initialized the genes of the model’s architecture using two approaches: (1) randomly distributing neurons across the three blocks, and (2) assigning all neurons to Dist, creating an initial architecture that was fully distributed, to test whether a structure with no initial modules at all can evolve into a modular structure. Under the same Face-Object dual task, we found that the face module was successfully formed in both initialization scenarios (Fig. S1), illustrating the robustness of the DAMP model to evolve a face module from various initial configurations.

The number of neurons gradually decreased in Dist and increased in M1 and M2, indicating a shift towards a modular structure. (C) The effect of sparseness in connectivity on modularity and accuracy. The x-axis represents the connectivity level. The red y-axis denotes the modularity score, measuring the degree of specialization in the network. The green y-axis represents the accuracy of the dual task. As the connections became less sparse (i.e., connectivity level approached 1), the model evolved into a more distributed structure, and the accuracy increased. Error bars indicate the standard error from 4 repeats.

One key parameter in this model is the connectivity level (i.e., sparseness), initially set to 0.5. To explore its effect on network architecture, we varied the connectivity level from 0.25 (highly sparse), 0.5, 0.75 to 1 (fully connected). We found that the modularity score decreased monotonically as connectivity changed from sparsely connected to densely connected (one-way ANOVA, *F*(3, 12) = 774.5, *p* < 0.001). Specifically, when the connectivity level increased from 0.5 to 1, 𝑁_𝐷𝐷𝐷𝑠𝑡_ increased, resulting in a more distributed network structure (Fig. 2C). That is, sparseness in connectivity is critical for the DAMP model to evolve a modular structure, as lower connectivity levels encourage the development of specialized modules by limiting the overlap in information processing between different blocks.

An interesting question is why the DAMP model evolves into a distributed structure with dense connectivity. To address this, we assessed the learning efficiency at different levels of connectivity. Specifically, we used the accuracy of Output1 and Output2 after 20 training epochs as an index of learning efficiency (Fig. 2C). We found that accuracy increased as connectivity changed from sparsely connected to densely connected (one-way ANOVA, *F*(3, 12) = 590.4, *p* < 0.001). This suggests that deleting neural connections impairs the model’s learning efficiency, leading the DAMP model, under the evolutionary algorithm, to evolve into a distributed structure in the absence of connectivity constraints. In other words, sparse connectivity is necessary for forming a modular structure.

Furthermore, models with sparse connectivity and a modular structure exhibit better learning efficiency than those with a distributed structure. To illustrate this, we compared the learning efficiency, indexed by the average number of training epochs required to achieve a certain accuracy (i.e., lower numbers indicating higher efficiency), between models with modular and distributed structures at a connectivity level of 0.5. Indeed, the learning efficiency for models with a modular structure was significantly higher than that for models with a distributed structure (39.3 versus 41.1 epochs, *t*(6) =-3.15, *p* = 0.020), suggesting that neural works with a modular structure are more efficient in fulfilling task demands when connections are sparse.

The sparseness of connectivity in biological neural networks is influenced by various factors, such as energy conservation (Tanaka et al., 2020; Wang et al., 2020). Previous studies have shown that significant deletion of connections greatly impaired network performance (Gale et al., 2019). The findings from the DAMP model suggest that modularity may arise as an adaptation strategy in response to challenges posed by sparse connections, rather than as an information processor specialized for certain stimuli or tasks. If this is the case, we would expect to identify modules for non-face objects in the DAMP model.

### Modularity emerged for non-face objects

To examine whether modules for non-face objects emerge automatically, we conducted two experiments by manipulating either task demands or stimulus type. In the first experiment, we compared faces versus non-face objects under the same task demand (e.g., face identification versus car identification, the Face-Car dual task). This experiment aimed to determine if modules would emerge independently of task demands. In the second experiment, we compared non-face objects under different task demands (e.g., car identification versus object categorization, the Car-Object dual task). This experiment was designed to assess if modules would form for processing non-face objects. These experiments allowed us to systematically assess the roles of task demands and stimulus types in driving the development of cognitive modules.

In both experiments, we observed similar patterns in the number of neurons across the three blocks (Fig. 3B & 3C), with the modularity scores approaching 1. Specifically, the modularity scores were 0.9996 ± 0.0011 for the Face-Car dual task, and 0.9989 ± 0.0014 for the Car-Object dual task (mean ± SD). This indicates that the number of neurons in Dist gradually decreased to near zero, leading to the emergence of a modular structure during evolution. In addition, the DAMP models processing other non-face objects (i.e., food and dog) also evolved into a modular structure (Fig. S2A & S2B). Taken together, these findings demonstrate that neither face stimuli nor identification tasks are necessary to form modular structures in the DAMP model. Rather, modular structures emerge as a general adaptation strategy in neural networks, driven by the constraints of sparse connectivity and the need for efficient information processing.

**Figure 3:**
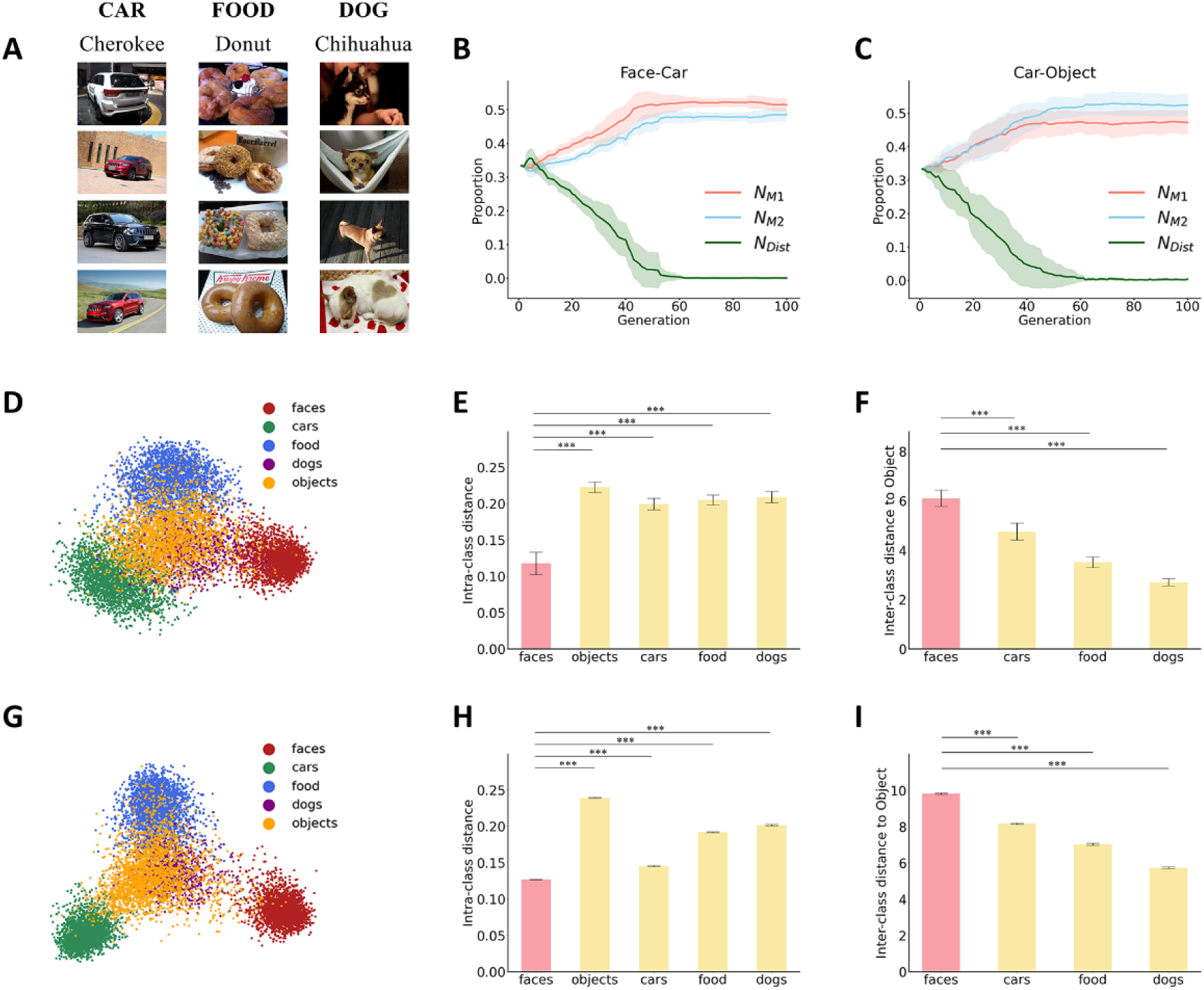
The specificity of the DAMP model. (A) Examples of stimuli from the Car/Food/Dog datasets. (B) Changes in number of neurons in the dual task of face identification versus car identification. The solid line represents changes in the number of neurons over the training period, with the shaded area indicating SD across 8 repeats. The colors indicate the different blocks (M1: red, Dist: green, and M2: blue). (C) Changes in the number of neurons in the dual task of car identification versus object categorization. (D) Two-dimensional visualizations of the point clouds of five stimulus categories in M1 (i.e., the face module). Each dot represents the neural representation for one exemplar, with colors indicating different categories (faces: red; cars: green; food: blue; dogs: purple; and daily objects: orange). (E) Intra-class distances of the five stimulus categories. The y-axis represents the variance of the point clouds, measuring the spread of neural representations within each category. (F) Inter-class distances between the centroid of the point cloud for daily objects and those for faces, cars, food, and dogs. The y-axis represents the distance in the high-dimensional neural space, indicating how distinct the neural representations of different categories are from daily objects. (G-I) The neural geometry, intra-class distances, and inter-class distances for the five stimulus categories in the neural space formed by the embedding vector of the DCNN encoder, which provide a comparison with the modular structure of the DAMP model. ***: *p* < 0.001

However, an apparent inconsistency exists between our findings and the fact that in the brain, the neural module for faces, but not for object categories tested here, is more easily identified (e.g., Grill-Spector & Weiner, 2014; Kanwisher, 2000). To address this inconsistency, we analyzed the neural representations in M1 where the face module was formed. To do this, we extracted the neural activations in M1 in response to all five categories (i.e., faces, daily objects, cars, food, and dogs) and then examined the neural geometry of the point cloud for each category. To visualize the geometry, we used principal component analysis (PCA) to project the high-dimensional neural geometry into a two-dimensional space. We found that the representation for faces was more tightly clustered and farther away from other stimuli (Fig. 3D, red). This observation was further quantified by the variances of the five point-clouds as an index for intra-class distances (Fig. 3E) (one-way ANOVA, *F*(1,4) = 131.4, *p* < 0.001). *Post-hoc* t-tests revealed that the intra-class distance of the point clouds for faces was significantly smaller than that of other categories (daily objects: *t*(14) = -16.2, *adjusted p* < 0.001; cars: *t*(14) = -12.3, *adjusted p* < 0.001; food: *t*(14) = -13.6, *adjusted p*  < 0.001; dogs: *t*(14) = -13.9, *adjusted p* < 0.001; Bonferroni corrected).

In addition, we used the point cloud of daily objects in the neural space as a reference and measured the inter-class distances between the centroid of the point cloud for daily objects and those for faces, cars, food, and dogs, respectively (Fig. 3F). One-way ANOVA revealed a significant difference in inter-class distances (*F*(1,3) = 210.0, *p* < 0.001). *Post-hoc* t-tests showed that the inter-class distance for faces was significantly larger than those for other categories (cars: *t*(14) = 7.5, *adjusted p* < 0.001; food: *t*(14) = 17.4, *adjusted p* < 0.001; dog: *t*(14) = 25.0, *adjusted p* < 0.001; Bonferroni corrected). Taken together, the neural geometry for faces in the face module was more tightly clustered and spatially more distant from that of daily objects. This geometrical characteristic enhances the distinctiveness of face category from others by concurrently narrowing and elevating neural tuning curves. Accordingly, the distinctiveness of the neural representations for faces may explain why the face module is more easily identified in the brain.

Interestingly, this characteristic in neural geometry apparently does not come from meta learning but rather from the pre-processing of the DCNN encoder that provides inputs to M1. For example, in M1 under the Car-Object dual tasks (i.e., car module), we found the same geometric characteristic for face representation – faces were more tightly clustered and farther away in the neural space (Fig. S2C) – even when faces were not present in the training dataset for the DAMP model. This finding is further confirmed by analyzing the neural geometry for faces in the DCNN encoder, where the same geometric characteristic for face representation was identified (Fig. 3G-I). This suggests that the distinctiveness of face representation likely comes from the inherent properties of faces (e.g., highly similar configurations and facial parts across individuals), making faces distinct from other object categories. However, the formation of cognitive modules is independent of stimulus properties and mainly relies on the structure constraints (i.e., sparseness of connectivity) to achieve better learning efficiency in the DAMP model. Thus, while face representations are distinct due to their stimulus properties, the emergence of modular structures in the DAMP model is driven by the model’s connectivity.

## Discussion

In this study, we used a computational modeling approach to investigate the origin of cognitive modules, focusing on the face module. We adopted the DAMP model, which combines a pre-trained DCNN encoder with a adaptable network using meta-learning to simulate the evolutionary process of forming specialized modules similar to the FFA.

We found that the DAMP Model autonomously evolved to form a specialized, modular structure, particularly a face module. This module emerged independent of stimulus type and task demands, suggesting that neither stimulus properties nor tasks are decisive factors in its development. Instead, the network’s structure, specifically the sparseness of connectivity, is critical. Sparser connectivity favors the formation of a modular structure to achieve higher learning efficiency, while denser connectivity leads to a more distributed structure. Therefore, our study reveals a fundamental property of neural networks with sparse connectivity: when optimizing to balance learning efficiency and energy conservation, these networks likely adopt a modular structure. This insight enhances our understanding of how cognitive modules, such as the face module, can naturally emerge in biological neural networks.

Methodologically, the DAMP model offers a novel perspective for understanding the origin of cognitive modules. By enabling neural networks to autonomously evolve and adapt without any prior network architecture, this approach mirrors evolutionary processes in biological systems. Traditional computational modeling studies often rely on pre-designed architectures to study the origin of cognitive modules (e.g., Blauch et al., 2022; Dobs et al., 2022), constraining their capacity to observe the spontaneous emergence of modular and distributed architectures. In contrast, our meta-learning framework retains the advantage of precisely controlling and manipulating influential factors while simulating evolutionary processes. By creating a competitive environment where network architectures evolve through training, selection, and genetic operations, the DAMP model allows the neural network to independently refine its structure and discover strategies with the best fitness for face processing. Thus, this meta-learning approach allows the exploration of a wide range of possibilities and the automatic evolution of structures based on information and tasks encountered, thereby providing a deeper understanding of how influential factors discovered by empirical studies drive the development of cognitive modules.

With this approach, we found that connectivity plays a crucial role in forming the face module, highlighting the close relationship between network structure and functional specialization. Previous studies have shown that connectivity significantly impacts neural functions, such as shaping dimensions via neuronal similarity (Kriegeskorte & Wei, 2021; Langdon et al., 2023), and resulting in the modularization and small-world nature of brain organization (Bassett & Bullmore, 2017; Sporns & Betzel, 2016). Specifically, the organization and specialization of the VTC are heavily influenced by connectivity among cortical areas within (Y. Zhang et al., 2024) and outside this region (Q. Chen et al., 2017; Stevens et al., 2015). Our findings reinforce this perspective, showing that the formation of the face module depends on sparse connectivity. Indeed, the reduction in connection costs leads to the emergence of more modular structures (Clune et al., 2013), which helps overcome the impairment in performance due to sparse connectivity. Therefore, with sparse connectivity, the DAMP model illustrates the emergence of the face module to balance the need to optimize learning efficiency and minimize connection costs (Y. Zhang et al., 2024), underscoring the critical role of network structure in driving functional specialization.

On one hand, our findings support the domain-specific perspective, positing that certain cognitive processes are uniquely tailored for specific content, separate from the neural circuits involved in processing others. On the other hand, our findings show that the DAMP model evolved into complete modularity regardless of tasks demands (identification versus categorization) and stimulus types (faces versus non-face objects), which appears inconsistent with the neuroimaging studies indicating that only a limited number of objects (e.g., faces, bodies, scenes) are processed in specialized areas in the human VTC (e.g., Downing et al., 2001; Epstein & Kanwisher, 1998). The analysis of the representational geometry of faces may reconcile this apparent inconsistency. One possible reason, as our model revealed, is the larger inter-class distances in the neural space between the point cloud for faces and those for non-face objects, signifying clear distinguishability. In addition, within the point cloud for faces, face exemplars are more tightly clustered, forming smaller intra-class distances than non-face objects, indicating more homogeneous representations. This clustered representation requires more nuanced distinctions between different face exemplars, facilitating efficient processing and recognition of faces. In fact, face perception is thought to have developed in primates and other vertebrates (Cuaya et al., 2016; Gross, 2008; Kendrick et al., 2001) due to heightened sensitivity to visual details (Leopold & Rhodes, 2010). Our findings, showing that the isolated and clustered face representation in the neural space was actually derived from the DCNN encoder rather than the evolutionary algorithm of meta-learning, align with this perspective. Therefore, the more readily formation of the FFA likely reflects the joint efforts of inherent structure (i.e., sparse connectivity) and external stimulus features (i.e., higher homogeneity within the face category).

While our study provides insights into the origin of cognitive modules, several limitations need future examination. First, our study highlights the critical role of connectivity in forming cognitive modules; however, the underlying mechanism remains unclear. One possibility is that connectivity modulates cognitive modularity by manipulating the dimensionality of neural geometry (Ma et al., 2023). Additionally, future studies should also explore other structural factors, such as neuronal response profiles (e.g., mixed selectivity: Bergoin et al., 2024; Cai et al., 2024; Ostojic & Fusi, 2024), on the modularity of neural networks. Second, our model, with its adaptable network architecture and genetic evolution algorithm, provides an evolutionary perspective on the origin of cognitive modules. However, this simulation is only at the algorithm level, with few biological constraints implemented. This lack of biological plausibility weakens the model’s explanatory power regarding the origin of cognitive modules in the human brain. Future studies should implement biological constraints in computational models to better reflect the nature of the human brain.

## Methods

### Stimuli and Tasks

To examine the impact of tasks and stimuli as two independent factors, we employed a diverse dataset comprising both coarse-grained and fine-grained object categories. We selected subsets from each dataset to ensure a balanced distribution of images across various object categories. Specifically, we utilized 500 facial identity images (100 images per identity) from the VGGFace2 database (Cao et al., 2018), 101 types of food (495 images per food type) from the Food101 database (Bossard et al., 2014), and 20,580 images representing 120 dog breeds (approximately 150 images per breed) from the Stanford Dogs database (Deng et al., 2009; A. Khosla et al., 2011). It is important to note that the dog images do not necessarily contain dog faces. From the CompCars database (Yang et al., 2015), we selected 500 cars, each containing 100 exemplars from various models and years. Thus, we constructed a database consisting of four object categories: faces, food, dogs, and cars, which is ready for the identification task. In addition, for the categorization task we selected 423 coarse-grained daily object categories (118 images per category) from the ILSVRC-2012 database (Deng et al., 2009). We manually excluded images related to the object categories designated for the identification task, thereby reducing the number of images for daily objects to 20,000.

All images were scaled to a minimum side length of 224 pixels, center-cropped, and normalized based on the mean and standard deviation values from ImageNet. To address the discrepancy arising from variations in the number of categories within the label of daily object (i.e., for the categorization task) and the number of exemplars within a category (i.e., for the identification task), we employed a more compact dataset for testing. Specifically, for each of the five stimulus categories—faces, daily objects, cars, food, and dogs—we randomly selected 100 identities (e.g., a person, a type of food, a dog breed, a car model, airplane from daily objects) from our database to ensure 40 exemplars for each identity. This method can effectively mitigate biases introduced by different numbers of images within the dataset.

We combined stimuli (faces versus non-face objects) and tasks (identification and categorization) into different types of tasks, which resulted in two types of tasks: face/car/food/dog identification task and object categorization task for daily objects. To form pairs of dual-tasks, we first adopted the most traditional dual-task: face identification versus object categorization to examine the emergence of the face module in the DAMP model. To examine the necessity of task demands, we matched the task demands in the dual task of face identification versus car identification, whereas to examine the necessity of face stimuli, we utilized the dual task of car identification versus object categorization. Finally, to demonstrate the robustness of our findings, we also included the dual tasks of food identification versus object categorization and dog identification versus object categorization.

### DAMP Model

To explore the origin of cognitive modules under the dual tasks, we employed the DAMP model. The DAMP model begins with a pre-trained convolutional image encoder, which vectorizes input images. Specifically, we utilized AlexNet (Krizhevsky et al., 2017) with frozen weights as the image encoder, and all pre-processed images were fed into it. To obtain image features not constrained to the 1000-category representations of ImageNet, we removed the final fully connected layer (4096-1000). Then, a sigmoid function was applied to normalize the 4096-dimensional features. Finally, we conducted a PCA on the entire dataset, retaining the first 1000 principal components, which accounted for approximately 80% of the explained variance.

The embedded features were then fully connected to three distinct hidden neuron blocks: M1, Dist, and M2. While maintaining the total number of neurons (𝑁_𝑡𝑜𝑡𝑎𝑙_), the distribution of neurons among the three blocks varies. These neurons operated in parallel and were all fully connected to the input embedding. These three blocks were connected to two output layers (Output1 and Output2), corresponding to two tasks, respectively. The outputs from M1 and Dist, activated by a sigmoid function, were concatenated and connected to Output1, while the outputs from M2 and Dist to Output2. That is, M1 and M2 were connected exclusively to Output1 and Output2, respectively, and Dist block was connected to both output layers. We suggest that different configurations of neurons in these three blocks represent various degrees of modularity, ranging from modular to distributed structures, with mixed structures falling in between. Accordingly, modularity is operationally defined by the modularity score, calculated as 1 − (𝑁_𝐷𝐷𝐷𝑠𝑡_/𝑁_𝑡𝑜𝑡𝑎𝑙_), which ranges from 0 to 1.

All neurons in the hidden blocks possess a non-linear sigmoid activation function. Each neuron in the output layer corresponds to a specific task, with the number of output neurons equal to the number of identities for that task, plus one additional neuron to indicate “other stimuli.” For instance, in the dual task of face identification versus object categorization, the activation of the additional neuron in Output1 signifies “not a face.”

The model was trained using the default Adam optimization algorithm with parameters 𝛽_1_ = 0.9, 𝛽_2_ = 0.999, and 𝜖𝜖 = 1𝑒^−8^ . The cross-entropy loss was calculated, and the weights were updated through backpropagation. Training was stopped after reaching up to 50 epochs or achieving an accuracy of 0.8, whichever came first, to expedite the process and minimize overfitting.

### Evolutionary algorithm of meta-learning

We utilized a genetic evolutionary algorithm to optimize the DAMP model’s architecture. Specifically, different neural network architectures were represented by different genotypes, and fitness was set as the direction of evolution. The goal was to either achieve a certain task accuracy faster or attain higher accuracy within a pre-set number of epochs. Genotypes also included learning parameters, as the optimal ones may vary with different architectures.

The selection process in the evolutionary algorithm affects the efficiency of evolution but rarely influences the final stable result. Therefore, following previous research, we chose the elite evolutionary algorithm to obtain a more stable evolutionary process. Specifically, we defined the fitness function as *fi𝑡𝑛𝑒𝑠𝑠* = 𝑒𝑝𝑜𝑐ℎ*_lim_* −𝑒𝑝𝑜𝑐ℎ + 𝑎𝑐𝑐, where 𝑒𝑝𝑜𝑐ℎ is the actual number of epochs required to reach the pre- set target accuracy within a limited number (𝑒𝑝𝑜𝑐ℎ*_li𝑚_* = 50), and 𝑎𝑐𝑐 is the accuracy at the last epoch. The accuracy is included because the model may not achieve the target accuracy within the pre-set maximum number of training epochs.

A gene in genotypes represents the specific expression of this gene individual. The evolved genotype, representing neural network architecture and learning parameters, comprises 14 genes, each a positive real number. Given that 𝑁_𝑡𝑜𝑡𝑎𝑙_ is fixed, 𝑁_𝑀1_ +𝑁*_Dist_* and 𝑁_𝑀1_ + 𝑁*_Dist_* are two free parameters rounded to integers from gene values. Each of the two layers (i.e., hidden and output) corresponds to a set of 12 = 2 × 2 × 3 parameters for training setup, including pairs of boundaries for learning rate, weight initialization, and bias initialization. Specifically, the remaining 12 gene loci are:

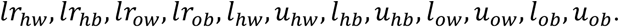

For the genotype population initialization step of the evolutionary algorithm, we generated 100 random 14-gene individuals, with the exception that two architecture parameters were set to 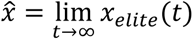 This ensured that all three hidden blocks had equal numbers, facilitating visualization and rapid simulation without impacting the final evolutionary outcomes. The trade-off in selecting the number of genotype populations and the maximum number of training epochs balances gene diversity and efficient evolution. Unlike classical genetic algorithms, where all genotypes evolve in parallel, we serialized the training of different models to simplify the process and avoid handling asynchrony, albeit at the expense of efficiency.

After one training epoch, the elite genotype with the highest fitness was selected and copied to fill one half of the total population by retaining the specific values of each locus unchanged. The remaining half underwent crossover to produce new individuals. For each locus, two values from two parents together provided a range for their offspring by specifying the upper and lower limit values. During this formation process, every genotype undergoes a certain degree of mutation by adding Gaussian noise to each locus. Note that noises on two loci for architecture were scaled since their values were much larger. The entire evolutionary process continued until a steady state was reached. That is, when the expression of the elite genotype no longer changed (differences compared to previous one are smaller than 1%) and survived. For each dual task, we repeated this process 8 times to ensure reliability of the results.

### Levels of connectivity in the DAMP model

To introduce biological constraints into the DAMP model, we implemented the connectivity level and examined how models differed in structures by setting the sparseness values to 0.25, 0.5, 0.75, and 1, respectively. In the experiments, to attain the model’s connectivity structure to the specified connectivity level, we randomly pruned connections between hidden and output neurons with proportion of 1 − 𝑓𝑓_𝐻𝐻𝐻_ by setting the weights of these chosen connections to 0.

To ensure a balanced and meaningful comparison, we adjusted the number of hidden neurons based on the connectivity level. For more connectivity, which risks overfitting or trivializing the task due to excessive connections, we reduced the number of hidden neurons. Conversely, for less connectivity, where too few connections might render the task unachievable, we increased the number of neurons. This modification ensured that, across all scenarios, the model underwent a reasonable number of epochs to achieve an accuracy rate close to 0.8.

Specifically, for the values of 0.25, 0.5, 0.75, and 1, we set 𝑁_𝑡𝑜𝑡𝑎𝑙_ to 1800, 900, 300 and 180, respectively. These numbers were chosen to ensure that the elite models in stable states of evolution would reach the target accuracy within at least 10 epochs. This way, the fitness of different network architectures varied significantly during evolution processes, allowing the evolutionary algorithm to efficiently select the elite one.

When assessing models’ accuracy, we fixed the total number of hidden neurons across different levels of connectivity. That is, we set 𝑁_𝑡𝑜𝑡𝑎𝑙_ to 900 across all levels and compared the accuracy that models could achieve in 20 epochs after 100 evolutionary generations. This approach ensured that 𝑁_𝑡𝑜𝑡𝑎𝑙_ remained the same, allowing us to compare their fitting capacity against different structures.

### Representational geometry of faces

We used the test dataset to evaluate the representations of stimuli within the neural space of hidden neurons across different models. These test images were processed through the elite model of the final generation, and their representations were examined in the neural space constructed by the neurons in M1. The dimension of this neural space equals the number of neurons, and thus an image is a vector of that dimension. In a typical high-dimensional neural space, each image is represented as a point, and images from a category form a point cloud. The centroid of each category was calculated as the average position of the points within that category, along with the variance of stimuli corresponding to each category.

We examined the geometry of point clouds in the original high-dimensional representational spaces. Thus, inter-class distances were calculated as the Euclidean distances between each pair of centroids of categories. We also calculated intra-class distances as the average of the variances of each category on all dimensions. To visualize the inter- and intra-class distances of categories, principal component analysis (PCA) was employed to reduce the high-dimensional neural spaces to two dimensions.

## Data availability

Data and code are available upon request by contacting the lead contact, Jia Liu.

## Contribution of authors

J.L. conceived the study. X.W. designed the experiments and prepared the stimuli. JR.L. developed the DAMP model and performed data analysis. X.W., JR.L., and J.L. wrote the paper.

## Acknowledgements

We would like to thank Mr. Jin Li for his insightful discussion and constructive advice on the evolutionary algorithm. This study was funded by the Shuimu Tsinghua Scholarship (X.W.), Beijing Municipal Science and Technology Commission & Administrative Commission of Zhongguancun Science Park (Z221100002722012), and Double First-Class initiative Funds for Discipline Construction.

## Declaration of conflicting interests

The author declared no potential conflicts of interest with respect to the research, authorship, and/or publication of this article.

## Notes

### Competing Interest Statement

The authors have declared no competing interest.

