## Supplementary material for "The Origin of Cognitive Modules for Face Processing: A Computational Evolutionary Perspective": Fig. S

### Supplementary Materials

**Figure S1. Robustness of the face module under the Face-Object dual task**. The genes of the model's architecture were initialized either by randomly distributing neurons across the three blocks (left panel) or by assigning all neurons to Dist, creating an initial architecture that was fully distributed (right panel). Under the Face-Object dual task, a modular structure was formed in both initialization scenarios.

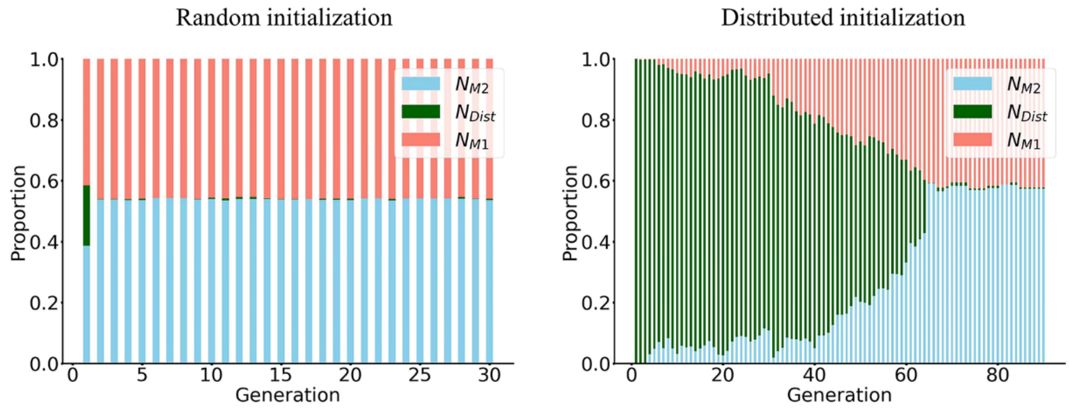


**Figure S2.** The emergence of cognitive modules for food (A) and dog (B) categories. Note that in the stable state, there were more neurons in M1 than M2, possibly due to the larger number of exemplars in the food and dog categories requiring more neurons for representation. (C) Representational geometry of neurons in M1 under the Car-Object dual task. Though face stimuli were not used for meta-learning in this task, an isolated and clustered face representation in the neural space was evident, suggesting that the characteristic of face representation likely derives from the DCNN decoder.

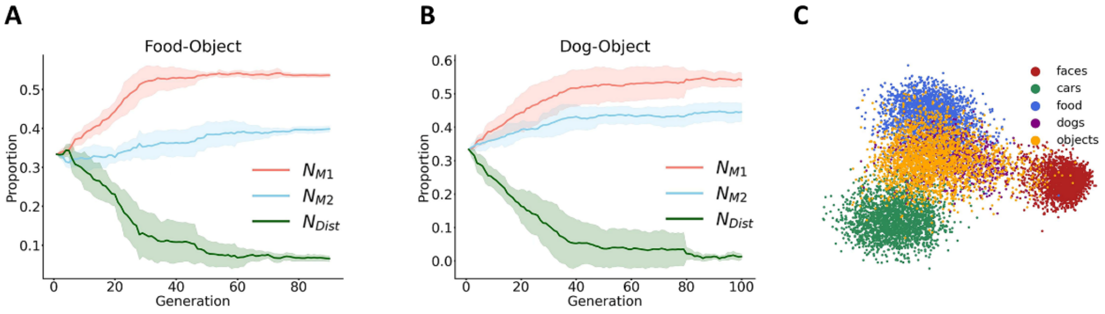
